# Programmable bacterial adhesion to plastic surfaces for enhanced biodegradation

**DOI:** 10.64898/2026.03.16.710745

**Authors:** Arianna Schneier, Benjamín O. Armijo-Galdames, Elizabeth C.H.T. Lau, Joanna C. Sadler

**Affiliations:** Institute of Quantitative Biology, Biochemistry and Biotechnology, School of Biological Sciences, University of Edinburgh, Roger Land Building, Alexander Crum Brown Road, King’s Buildings, Edinburgh, EH9 3FF, UK

**Keywords:** plastic, engineered adhesion, curli, antigen 43, PET degradation, co-expression

## Abstract

Colonisation of plastic surfaces by microbial biofilms offers a promising starting point for engineering efficient biodegradation systems. However, most studies to date focus on characterisation or prevention of biofilms on plastics in diverse environments and the potential biotechnological application for these systems has been underexplored. To address this, we report the efficient adhesion of *Escherichia coli* cells to a range of plastic surfaces through overexpression of two key determinants of bacterial biofilm formation; curli and Antigen 43 (Ag43). A general trend of higher total biomass was observed from curli-mediated adhesion, but more uniform adhesion from Ag43 overexpression. We further demonstrate application of this technology through inducible adhesion of *E. coli* to polyethylene terephthalate (PET) surfaces and concurrent secretion of the PET depolymerase PHL7. Co-overexpression of curli fibres and secreted PHL7 resulted in 5.6-fold increase in terephthalic acid release in comparison to the non-adherent control. These methods offer a general approach to programmable adhesion of genetically tractable cells to plastic surfaces and concurrent secretion of degradative enzymes, and are anticipated to be broadly applicable across the field of plastic bioremediation technologies.

## Introduction

The prevalence of plastic in natural environments has triggered a surge of interest in developing degradation methods to remove recalcitrant polymers from circulation. These include chemical and biotechnological processes for both single polymer types and, more recently, mixed plastic waste^1–3^. However, challenges remain for plastic degradation to be efficient, robust and low-cost enough to be implemented at scale. In particular, most methods to date rely on elevated reaction temperatures, high pressures, use of fossil-fuel derived reagents and solvents or in the case of biocatalysed processes, high loadings of purified enzyme. Therefore, methods to improve the efficiency and decrease costs of these processes remain an important research focus.

A fundamental challenge in plastic depolymerisation is the heterogeneity of reaction mixtures. In the context of biocatalysed degradation, adsorption of the biocatalyst onto the plastic surface is a key determinant of degradation rate^4,5^. Indeed, studies have shown that approaches to localise plastic-degrading enzymes at the plastic-solvent interface significantly improves degradation efficiency^6,7^. For example, Zhu and co-workers applied a Biofilm-Integrated Nanofiber Display (BIND) platform to generate a whole-cell biocatalyst comprising surface-displayed PETase via CsgA, the monomeric repeating subunit of curli fibres, for degradation of PET microplastics^8^. Though curli is well known for its involvement in adhesion to surfaces^9–11^, the interaction between BIND-PETase and the plastic substrate was not characterised, though addition of 0.02% sodium tetradecyl sulphate led to an increase in the presence of degradation products and was hypothesised to improve interaction between PET and PETase^12^. Other strategies have investigated the pairing of surface-displayed hydrophobins^13,14^ or adhesive proteins with PET hydrolases^15^ for improved localisation, all of which exhibited higher PET-degrading activity than either purified PETase or the surface-displayed PET hydrolase alone.

Intriguingly, in the relatively brief time frame in which plastic pollution has existed, Nature has already begun to deploy biological processes for the degradation and even valorisation of abiotic polymers. Many of these systems are mediated by microbial communities which colonise plastic surfaces, forming biofilms that create favourable microenvironments for secretion and activity of degradative enzymes directly at the microbe-polymer interface^16–21^. Further, biotechnologists have demonstrated that augmenting these naturally occurring biofilm communities can accelerate plastic degradation. For example, Syranidou and co-workers bioaugmented native marine consortium for degradation of weathered PE films, showing biofilm-treated samples to give a 4.5-fold increase in percentage weight loss relative to indigenous consortia over a 6-month period^22^.

The first step in these processes is localisation of microbial cells to the plastic surface. Surface association is mediated by extracellular, cell surface-associated adhesion factors singly or in combination. Flagella, type I fimbriae, and curli are involved in initial surface adhesion, while the production of a colanic acid rich exopolysaccharide (EPS) matrix and adhesins such as Antigen 43 (Ag43) and pili aid biofilm maturation^23–27^. Curli, Ag43, and EPS in particular have all been demonstrated as programmable platforms for cell-surface display of functional cargos^12,28,29^ yet have been underexploited in the context of mediating programmable association with plastic surfaces. Of particular interest to our study were Ag43 and curli. Ag43 is a natively expressed via expression of the *flu* gene in *E. coli* as a self-recognising adhesin, protruding ~10 nm from the outer cell membrane. Two Ag43 units from neighbouring cells form a ‘Velcro’-like interaction mediated by salt bridges and hydrogen bonding at the protein-protein interface, inducing microcolony formation^30–32^. Meanwhile, curli fibres form via a self-assembly of secreted CsgA monomers on the outer-membrane bound CsgGFB complex to form rigid, 2-5 nm long amyloid fibres around the cell^9^. Native curli expression is controlled by a complex regulatory network which can be upregulated under varied conditions including conditions of pH, temperature, abiotic stress and nutrient limitation^10,33,34^. Despite the roles of these proteins in biofilm formation being well studied, their use in a whole-cell engineering for plastics bioremediation remains limited. A study by Leech and coworkers found that *Escherichia coli* strain PHL644, which overproduces curli, showed better adhesion to hydrophobic surfaces such as polycarbonate compared to polystyrene and polytetrafluorethylene (PTFE) compared to glass. Growth on PTFE was subsequently optimised by changing the glucose concentration, temperature and shaking, amongst other conditions^35^. In a separate study, Howard and McCarthy demonstrated that pairing the overexpression of diguanylate cyclases, which promote biofilm formation, with novel polyester-degrading enzymes resulted in a ~1.4-fold increase in percentage of polycaprolactone weight loss compared to the depolymerase enzymes alone^4^. Building on these studies, we hypothesised that biofilm-mediating adhesins could be deployed for programmable adhesion to plastic surfaces, and further a biomimetic approach pairing secretion of a depolymerase could improve degradation efficiency (Figure 1).

**Figure 1.**
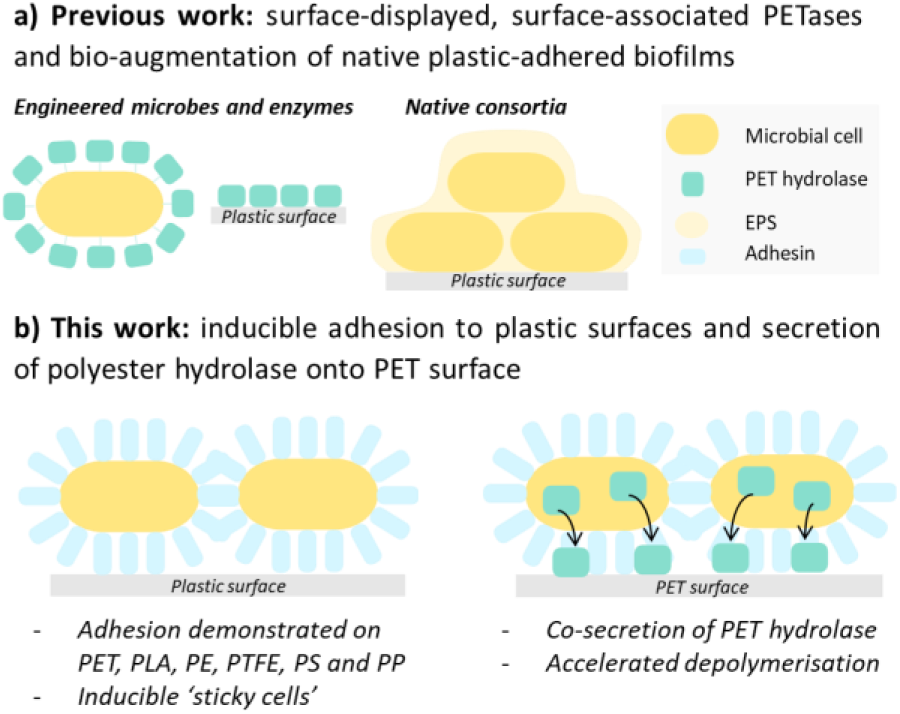
(a) Previous work on enzyme-PET co-localisation and microbial attachment to plastic surfaces has focussed on surface displayed enzymes, use of hydrophobic fusion peptides and proteins to localise at surface, and study of unmodified biofilms on abiotic surfaces. (b) This work presents an inducible platform for controlled microbial adhesion to varied plastic, and demonstration of an application in accelerated biodegradation through coupling with a secreted PET hydrolase.

Herein, we report the use of Antigen 43 (Ag43) and curli for inducible adhesion of *E. coli* cells to a range of commercial standard and post-consumer plastic surfaces. We further show potential application of these ‘sticky cells’ for accelerated biodegradation of an exemplar plastic, poly(ethylene terephthalate) (PET), demonstrating this as a versatile method for microbe-plastic co-localisation and a novel approach to accelerating biodegradation of PET.

## Materials and methods

### General procedure for Ag43 or curli-mediated plastic adhesion

Plastic samples (Table S7) were cut into 1-cm squares, washed with 70% ethanol and dried overnight at 37 °C. Microbial growth on plastic surfaces was achieved by modifying a previous protocol^36^. Briefly, overnight cultures were incubated in Lysogeny Broth (LB) media containing 100 μg/mL ampicillin and used to inoculate (1% v/v inoculum) sub-cultures in LB media containing 100 μg/mL ampicillin. The cultures were incubated at 30 °C until OD_600_=0.5-0.9 was reached and then used to inoculate assay plates (24-well non-tissue culture treated; Nunc, 144530 or Corning Costar, CLS3738). One sterilised plastic square was added per well for assessing adhesion to different plastic types. 1 mL of M63 media (Table S8) containing 100 μg/mL ampicillin was added to each well and inoculum described above (1% v/v final volume). For induced wells, arabinose was added to a final concentration of 0.1% w/v. Plates were sealed with gas permeable membranes (Breathe-Easy, Diversified Biotech BEM-1) and incubated at 30 °C with rocking (VWR incubating rocker) or orbital shaking (ThermoSTAR, 230 rpm, 0.7 mm orbit radius) for 18-28 hours.

### Quantification of biomass adhesion with crystal violet

A crystal violet (CV) assay was used to quantify the adhesion of biomass to plastic surfaces. At the end of the biofilm adhesion assay, supernatant was carefully removed from plates by pipetting. Wells were washed three times with 1 mL sterile water, before adding 500 μL 0.1% w/v aqueous crystal violet solution (Alfa Aesar). Wells were then gently washed three times with 1 mL sterile water. If the adhesion assay was performed on plastic squares, the plastic squares were then transferred to a fresh plate. The squares or wells were left to dry at 37 °C overnight then photographed to record the distribution of the stain across the wells or squares.

To quantify crystal violet adhesion to squares, 30% w/v aqueous acetic acid was added to each well and 200 μL transferred to a 96-well plate. Absorbance at 550 nm was measured using a SPECTROstar plate reader and blank corrected with 30% aqueous w/v acetic acid.

### General procedure for pairing of Ag43 and curli-mediated adhesion with enzyme secretion

Overnight cultures of the relevant strain(s) were grown in LB media containing 100 μg/mL ampicillin and 50 μg/mL kanamycin and used to inoculate (1% v/v inoculum) sub-cultures in LB media containing 100 μg/mL ampicillin and 50 μg/mL kanamycin. The cultures were incubated at 30 °C until OD_600_=0.5-1.3 and then used to inoculate (1% v/v inoculum) M63 media with 100 μg/mL ampicillin, 50 μg/mL kanamycin and 0.1% arabinose in either assay plates or 5 mL Corning tubes. Cultures were grown at 30 °C or 37 °C as specified for 5 hours, prior to addition of isopropyl β-D-1-thiogalactopyranoside (IPTG, 0.1 mM final concentration) and incubation as indicated.

### Hydrolase activity assay

The *p*-nitrophenyl butyrate (pNPB, Sigma-Aldrich) assay was used to quantify the activity of PET hydrolases through a colorimetric change when pNPB is hydrolysed to *p*-nitrophenol^12^. The pNPB reaction media comprised phosphate-buffered saline (PBS) containing 1 mM pNPB and 5% v/v dimethyl sulfoxide, which was used to solubilise the pNPB prior to addition to PBS.

This assay was used to determine hydrolytic activity of bacteria adhered to the plastic surfaces, and of extracellular secreted fractions. For screening the supernatant, culture was transferred to a fresh Eppendorf tube and clarified by centrifugation (5 minutes, 15800 x g), and 5 μL of the clarified supernatant added to 95 μL pNPB reaction mixture in a 96-well plate (Greiner F-bottom). To determine activity of cells adhered to plastic, the general procedure for mediated adhesion was applied, then followed by the removal of supernatant through careful pipetting. Plastic squares were washed three times with 1 mL sterile water before transferring to a fresh 24-well plate and adding 570 μL pNPB solution (as above) and 30 μL PBS to each well.

All pNPB assay plates were incubated for 30 minutes at 30 °C with rocking (20 speed, 7 tilt), prior to measuring absorbance of 100 µL samples at 405 nm using a SPECTROstar plate reader. All experiments were blank corrected with unreacted pNPB solution, and NaOH mediated pNPB hydrolysis was used as a positive control.

### Adhesion-accelerated biodegradation of PET

PET flakes (cryo-milled^37^, 2 mm average particle size, kindly provided by Prof. A. Pickford, University of Portsmouth) were dispensed into 5 mL screw cap tubes (Corning, AXYSCT10MLS) with 5-15 mg per tube. Plastic-adhered cells were prepared according to the general procedure above, with the modification that cultures were inoculated in 5 mL Corning tubes instead of plates. Cultures were induced first with arabinose (0.1% arabinose final concentration) at the time of tube inoculation, incubated with shaking (orbital or linear) for 5 hours at 30 °C and then further induced with IPTG (0.1 mM or 0.45 mM final concentration, as indicated). Plastic-containing cultures were then incubated at 30 or 37 °C with shaking (orbital or linear, as indicated) for 7 days, prior to centrifuging (5 minutes, 15800 x g) and analysis of the supernatant by HPLC (see Supplementary Methods). Statistical analysis was conducted using GraphPad Prism. The normality was assessed using the Shapiro-Wilk test. The comparisons were assessed using an unpaired t-test with Welch correction and *p*-value < 0.05 were deemed significant (denoted as *).

## Results and Discussion

We focussed our study on the use of curli and Ag43 for adhesion of *E. coli* cells to plastic surfaces, both of which have indirectly been described as mediating adhesion to plastic surfaces, although these interactions have been poorly quantified for different plastic types^35,36,38^. To determine their ability to confer adhesion to plastic, *E. coli* MG1655 was transformed with an arabinose inducible plasmid encoding overexpression of Ag43 (pAg43) or curli (pCurli) and adhesion to untreated polystyrene (PS) 24-well plates was analysed in the absence and presence of arabinose (Figure 2). Visual analysis of crystal violet staining clearly showed increased adhesion when curli or Ag43 were overexpressed, with more homogeneous coverage observed for Ag43 (Figure 2a). Quantification of crystal violet staining confirmed a 2.6-fold increase in biomass adhesion for curli, but only 1.3-fold increase for Ag43. To interrogate the extent to which native biofilm formation was contributing to these data and to generate a chassis for tightly controlled plastic adhesion, we generated single and double knock-out (KO) strains of *E. coli* MG1655 deficient in the native Ag43 gene (denoted *flu*) and/or *csgA*, the repeating external subunit of curli fibres^39^. An increase in biomass adhesion was observed for all KO strains, demonstrating an inducible system for microbe-PS adhesion (Figure 2a, Figure 2b and Figure S1). Interestingly, all three KO strains produced higher adhesion, 11.7-fold increase for *E. coli* MG1655Δ*flu*, 8.6-fold increase for *E. coli* MG1655Δ*csgA*, 9.3-fold increase for *E. coli* MG1655Δ*csgA*Δ*flu*, upon induction of pCurli than the parent strain, suggesting that native transcription or translation of the *csgA* or *flu* may inhibit high expression levels of these proteins. Less background adhesion was observed for MG1655Δ*csgA*Δ*flu* (uninduced) than the parent strain carrying pAg43, whereas background adhesion was lower in both cases for strains carrying pCurli. Together, these data demonstrate *E. coli* MG1655Δ*csgA*Δ*flu* as a versatile chassis for generation of an inducible platform for plastic adhesion (Figure 2b).

**Figure 2.**
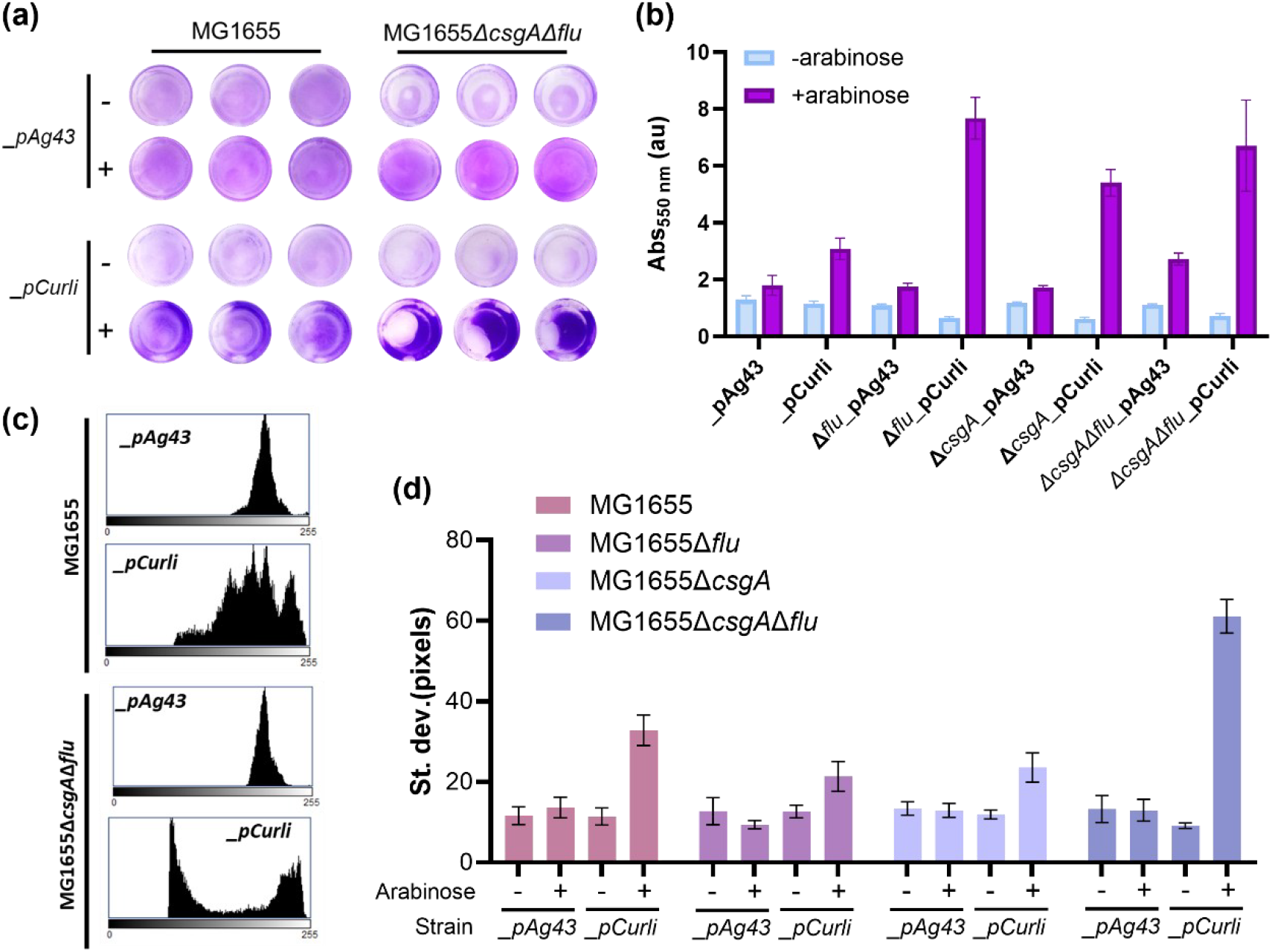
Inducible adhesion of E. coli to PS surfaces. (a) Crystal violet staining of total biomass adhesion of induced and non-induced MG1655 and MG1655ΔcsgAΔflu harbouring pAg43 or pCurli; (b) Quantitative analysis of crystal violet staining of induced and uninduced parent, single and double KO strains harbouring pAg43 or pCurli. (c) Representative histograms from image analysis of stained PS wells with induced samples showing degree of heterogeneity of coverage (full dataset in Figure S2). (d) Standard deviation from the mean image intensity across PS well for induced and uninduced parent, single and double knockout strains harbouring pAg43 or pCurli. Adhesion assay conditions: 0.1% arabinose, 30 °C, 230 rpm, 20 h.

To further interrogate the coverage of plastic surfaces by our engineered strains, we performed quantitative image analysis on the well-images shown in Figure 2a. This showed that with and without induction pAg43 has a similar intensity and distribution, whilst pCurli has greater standard deviation after induction indicating a greater degree of heterogeneity of distribution of adhesion across the surface (Figure 2c, Figure 2d, Table S9 and Figure S2).

Encouraged by these data, we next set out to assess whether this approach could be used as a general approach for adhesion of microbial cells to a range of plastic surfaces. We performed crystal violet staining assays for adhesion to poly(ethylene terephthalate) (PET), poly(lactic acid) (PLA), polypropylene (PP), poly(tetrafluoroethylene) (PTFE) and low-density poly(ethylene) (LDPE), representing both hydrolysable and non-hydrolysable plastics predicted to have varying surface properties. Experiments were performed on commercial grade substrates (Figure 3a-b) to enable standardisation across the field, as well as four authentic post-consumer plastic samples (PET, PLA, PP and LDPE, Figure 3c, Figure S3) to explore the application of this method to real-world waste streams. The strains overexpressing pAg43 exhibited similar adhesion profiles for both the commercial and post-consumer plastics, while pCurli adhered less to post-consumer plastics compared to the commercial samples. Whilst we hypothesised that adhesion may correlate to surface hydrophobicity, measured water contact angle measurements (Figure S4) did not correlate with observed adhesion, corroborating previous studies demonstrating complex relationship with other surface properties including surface charge and topology^25,40,41^.

**Figure 3.**
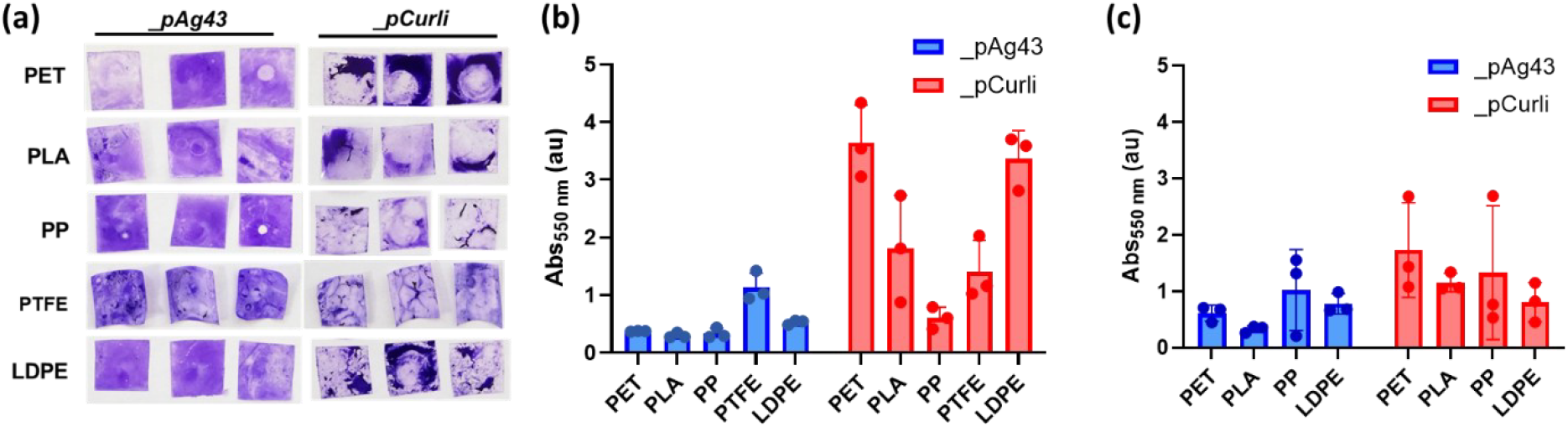
Adhesion of E. coli MG1655ΔcsgAΔflu expressing _pAg43 or _pCurli to different plastics. (a) Crystal violet imaging of commercial plastic-adhered biomass across the plastic surface and quantification of (b) commercial and (c) post-consumer plastic-microbial adhesion. Adhesion assay conditions: 0.1% arabinose, 30 °C, 230 rpm, 20 h.

We anticipate these plastic-adhering strains to have a range of applications across the field of plastic bioremediation and whole cell biocatalysis. One application we were keen to explore was use of surface-adhered cells for biomimetic plastic degradation. Specifically, we hypothesised that localisation of microbes to the cell surface could accelerate the rate of biodegradation of the plastic by a suitable secreted hydrolytic enzyme (Figure 4a). To test this hypothesis, we assembled plasmids encoding secretion of PET hydrolases via *N*-terminal fusion of the B1PelB secretion tag^42^. Using MG1655Δ*csgA*Δ*flu* as the chassis, protein production of both *Is*PETase^S238F/W159H43^ and PHL7^44^ was confirmed through SDS-PAGE (Figure S5a). Hydrolytic activity of the culture supernatant was assessed using the *p-*nitrophenyl butyrate (pNPB) assay, which showed a 2.4-fold higher activity for PHL7 than *Is*PETase^S238F/W159H^ (Figure S5b). PHL7 was therefore selected for further experiments. Culturing conditions for sequential expression of Ag43 or curli then PHL7 were optimised, with the key determining features identified to be a 5-hour delay between arabinose-mediated induction of Ag43 or curli prior to induction of PHL7 secretion with 0.1 mM IPTG (Figure S6-7).

**Figure 4:**
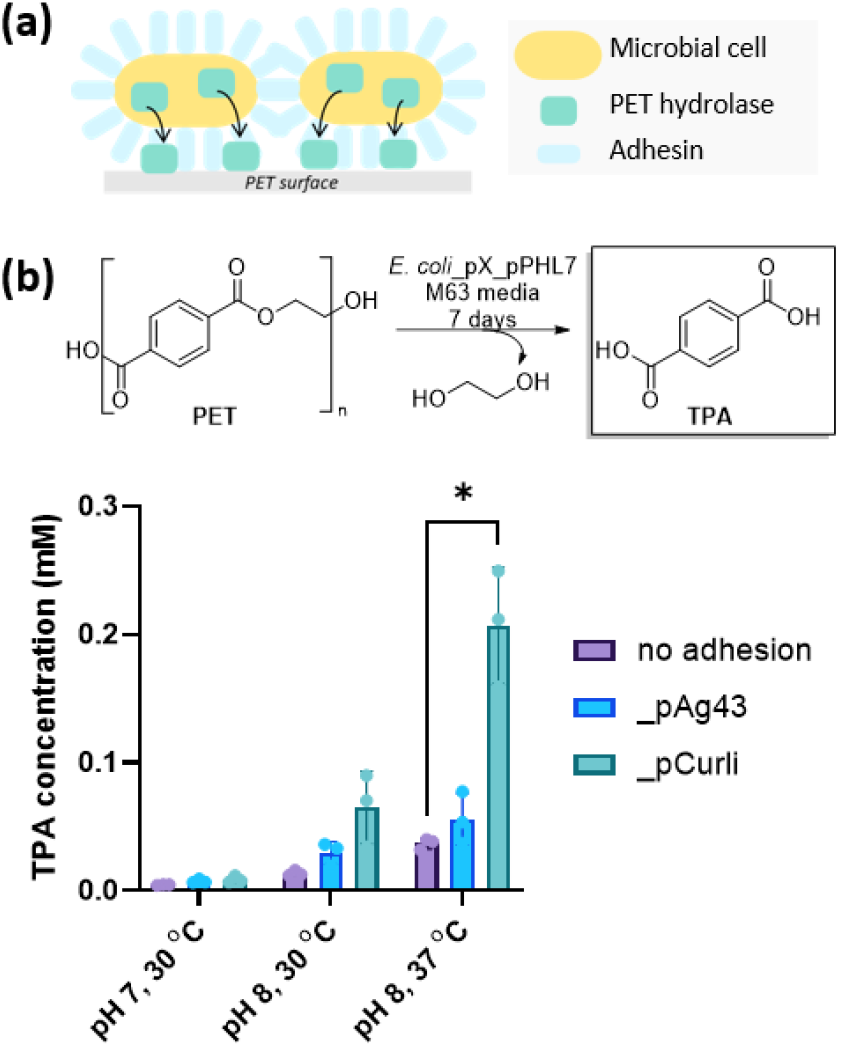
Accelerated PET biodegradation through co-expression of surface adhesins. (a) Schematic of overall process, in which PET hydrolyse is secreted directly to the PET surface; (b) HPLC analysis of TPA release from PET substrate through co-expression of PHL7 and Ag43 or curli (*: p < 0.05, unpaired t-test with Welch correction). Degradation assay conditions: 5 h induction gap, 0.1% arabinose, 0.1 mM IPTG, 7 days

We next sought to determine whether pairing over-expression Ag43 or curli with PHL7 increased PET hydrolysis efficiency. PET degradation efficiency was determined by detection of terephthalic acid (TPA) in the culture supernatant 7 days after PHL7 expression, with all experiments using the chassis strain MG1655Δ*csgA*Δ*flu* to suppress native biofilm formation. Whilst negligible PET degradation was detected for all strains at pH7, at pH8, comparison of TPA concentrations from a strain lacking enhanced adhesion (pBAD empty vector control) and strains harbouring pAg43 or pCurli all demonstrated a small increase in TPA concentrations for PET-adhered strains. At pH 8 and 30 °C, co-expression of Ag43 and curli resulted in a 2.2-fold and 4.8-fold increase, respectively, in TPA concentration after 7 days (*p*=0.0778 and 0.0875). However, the greatest improvement was observed at pH8 and 37 °C, where a 5.6-fold, (*p*=0.0223) increase in TPA concentration was measured for the pCurli strain. These data support our hypothesis that co-expression of microbial adhesins can enhance plastic biodegradation by a secreted hydrolase, relative to controls lacking surface adhesion (Figure 4b). This study is one of the first examples of using a synthetic biology approach to biomimetic plastic degradation using surface co-localisation paired with depolymerase secretion, suggesting this could be a useful tool for enhancing engineered plastic biodegradation systems.

## Conclusions

This study systematically explores the use of two surface-displayed proteins, Ag43 and curli fibres, for controllable adhesion of *E. coli* cells to plastic surfaces. We demonstrate that both proteins effectively mediate microbial adhesion to a wide range of commercial and post-consumer plastic types and note key differences in the morphology of surface adhesion. Specifically, Ag43 consistently formed more uniform, less dense layers across the plastic surface, with less total biomass but higher surface coverage, whereas curli typically forms localised patches of densely aggregated biomass on the surfaces, leading to higher variability across samples. It is anticipated that this technology could have diverse applications across the field of biotechnology, such as adherence of whole-cell biocatalysts to solid supports in flow biocatalysis applications, for example. In this study, we used the system to mimic nature’s newly evolved method of plastic biodegradation, namely adhering cells to a plastic surface and secreting a degradative enzyme, which was shown to significantly increase monomer release relative to controls lacking adhesion. This approach demonstrates how synthetic biology can leverage and mimic natural microbial mechanisms to accelerate degradative processes and is anticipated to have general utility across the field of plastic biodegradation.

## Acknowledgements

JCS acknowledges as a BBSRC Discovery Fellowship (BB/S010629/1); JCS and ECHT acknowledge the Preventing Plastic Pollution with Engineering Biology (P3EB) Hub (BB/Y007972/1). AS acknowledges the UKRI Biotechnology and Biological Sciences Research Council (BBSRC) grant number BB/T00875X/1. BAG acknowledges Darwin trust of Edinburgh PhD studentship. JS and AS are grateful to Prof. Andrew Pickford for provision of cryo-milled PET for use in degradation assays.

