## Supplementary Materials for "Programmable bacterial adhesion to plastic surfaces for enhanced biodegradation"

### Supplementary Information

**Contents**

1 General materials and methods ... 2

1.1 Molecular biology techniques ... 2

1.2 Analysis of TPA release from PET degradation assays ... 2

1.3 ImageJ analysis ... 3

1.4 Protein expression and SDS-PAGE ... 3

2 Table of strains and plasmids ... 3

2.1 Gene Sequences ... 3

2.2 Plasmid assembly ... 5

2.3 Strains ... 7

2.3.1 Knockout strain generation ... 7

3 Plastic substrates ... 8

4 Additional experimental data ... 9

4.1.1 Media recipe ... 9

4.1.2 Crystal violet staining and ImageJ analysis of adhesion to PS plates ... 10

4.1.3 Engineered adhesion to post-consumer plastics ... 13

4.1.4 Water contact angle measurements ... 14

4.1.5 Expression and activity of PET hydrolases ... 14

4.1.6 Optimisation of pairing adhesion and secretion ... 15

5 References ... 16

### 1 General materials and methods

Unless otherwise stated, reagents and high-performance liquid chromatography (HPLC) solvents were purchased from commercial suppliers and used without prior purification. All experiments involving microbial cultures were performed as independent biological replicates. The following abbreviations are used throughout: Antigen 43 (Ag43), low-density poly(ethylene) (LDPE), isopropyl β-D-1-thiogalactopyranoside (IPTG), Lysogeny Broth (LB), polyester hydrolase Leipzig 7 (PHL7), polymerase chain reaction (PCR), poly(ethylene terephthalate) (PET), poly(lactic acid) (PLA), poly(propylene) (PP), poly(styrene) (PS), poly(tetrafluoroethylene) (PTFE).

#### 1.1 Molecular biology techniques

Unless otherwise stated, all genes used in this study were ordered as lyophilised double stranded DNA (Twist Biosciences™). Single strand oligonucleotide primers were synthesised by Integrated DNA Technologies (IDT).

Synthetised genes were resuspended in MilliQ water to a final concentration of 10 ng/µL. PCRs were conducted in a 50 µL using Q5® High-Fidelity DNA Polymerase (New England Biolabs, NEB) with 0.1 ng DNA template and an extension time of 30 seconds per kb. The resulting products were either purified using a Qiagen™ QIAquick Gel Extraction Kit or treated with DpnI (NEB) for a 1 hour at 37 °C, deactivated at 80 °C for 20 minutes, then purified using the Qiagen™ QIAquick PCR purification kit according to the supplier's protocol. FastDigest™ (ThermoFisher) restriction enzymes were used for cloning. Digestion reactions were performed at 20 µL scale, containing 1 µg of purified DNA and incubated at 37 °C for 1 hour. Reaction products were purified with Qiagen™ gel extraction kit according to the supplier's protocol.

DNA ligations were performed using T4 ligase (NEB or Promega) and BsaI-HF®v2 or BsmBI-v2 (NEB) mediated cleavage. Colony PCRs were carried out using OneTaq® Quick-Load® 2X Master Mix (NEB) with 25 µL reaction scale and an extension time of 60 seconds per kb.

The ligated plasmids (5 µL) were used to transform chemically competent *E. coli* DH5α, MG1655, MG1655Δ*csgA*, MG1655Δ*flu*, or MG1655Δ*csgA*Δ*flu* (as stated) cells by heat-shock at 42 °C for 42 seconds, followed by addition of 1 mL SOC media and incubation at 37 °C for 1 hour (1 plasmid) or 1.5 hours (2 plasmids) at 600 rpm (Eppendorf ThermoMixer C). Transformants were selected via plating onto LB agar containing the appropriate antibiotic(s) (ampicillin at 100 µg/mL and/or kanamycin at 50 µg/mL) and incubated overnight at 37 °C. Successful uptake of the desired constructs was confirmed by colony PCR, plasmids were isolated using Qiagen™ QIAprep Spin Miniprep Kit, and sequences were verified by Sanger sequencing (GENEWIZ from Azenta Life Sciences) or full plasmid sequencing (Plasmidsaurus).

For agarose gels, 1% w/v agarose TAE gels (0.1% v/v SYBRSafe™ (Invitrogen)) were run at 100 V for 45-60 minutes. For SDS-PAGE, NuPAGE™ 4-12% Bis-Tris gels (Invitrogen) were run for 1 hour at 200 V in 1xMOPS buffer, before staining with InstantBlue® Coomassie Protein Stain (Abcam Plc) or Coomassie stain (0.1% Coomassie Brilliant Blue R-250, 40% v/v ethanol, 10% v/v acetic acid). Gels were visualised using a Gel Doc.

#### 1.2 Analysis of TPA release from PET degradation assays

Degradation assay supernatants were analysed by High Performance Liquid Chromatography (HPLC) using a Vanquish Core HPLC instrument equipped with a ZORBAX RRHT StableBond C18 Column (3.0 x 50 mm, 1.8 µm, 600 bar pressure limit). To a polypropylene 96-well plate (Greiner, 651201), 100 µL of sample was added to 50 µL dimethylformamide, 15 µL aqueous 1 mM solution of zingerone

(Fluorochem, 464499) in 1 M HCl. If the pH of the sample was 1-2, the sample was topped up to 200 µL with MilliQ-water. If the pH of the sample was >2, aqueous 1 M HCl was added to the sample until the correct pH was reached and then the sample was topped up to 200 µL with MilliQ-water. The plate was then sealed with a pierceable film (Zone-Free) and 10 µL was injected per sample for analysis.

The column temperature was set at 20 °C. All analytes were detected at 250 nm, 280 nm, 240 nm and 210 nm by a UV-vis detector and then quantified by an external calibration curve relative to zingerone. The gradient HPLC was used with 0.4 mL/min flow rate mixing water with 0.1% trifluoroacetic acid (TFA) (Solvent A) and acetonitrile with 0.1% TFA (Solvent B) (Table S1).

**Table S1.** Gradient HPLC make up of solvent B (acetonitrile with 0.1% TFA) compared to solvent A (water with 0.1% TFA).

| Time (min) | %B |
|---|---|
| **0 to 1.5** | 5 |
| **1.5 to 14** | 5 to 30 |
| **14 to 15** | 30 to 5 |
| **15 to 20** | 5 |

##### 1.3 ImageJ analysis

Images were compiled and imported into ImageJ (Fiji) 1.54p[1]. Images were converted to 8-bit grayscale and the oval selection tool was used to select the discs with adhered cells. The histogram tool under Analyze was then used to generate histograms (0-255) with count, mean, standard deviation, mode, intensity of mode, max and min reported.

##### 1.4 Protein expression and SDS-PAGE

For the screening of PET hydrolases on their own, overnight cultures in LB broth containing 50 µg/mL kanamycin were used to inoculate (1% v/v inoculum) sub-cultures. Sub-cultures were incubated (30 °C, 200 rpm) in LB media (containing 50 µg/mL kanamycin) until an $OD_{600}$=0.5-0.8 was reached and a 500 µL sample was taken for analysis of pre-induction protein expression ('uninduced'). Cultures were put on ice for 20-30 minutes after which heterologous protein expression was then induced by addition of isopropyl β-D-1-thiogalactopyranoside (IPTG, 0.45 mM final concentration). The cultures were then incubated at 20 °C and 200 rpm overnight. The following day, the final $OD_{600}$ was measured and end-point samples were taken. For the supernatant (secreted fraction), undiluted samples were clarified by centrifugation and 10 µL of the supernatant was removed for analysis. For the pellet (total protein), the culture was diluted to the same $OD_{600}$ as the uninduced sample, pelleted by centrifugation and then the pellet was resuspended to 10 µL with MilliQ-water. Samples were then prepared for SDS-PAGE according to the standard procedure, loading 10 µL per well of a pre-cast gel (NuPAGE™ 4-12% Bis-Tris). For the pNPB assay, the hydrolase activity assay protocol was followed.

### 2 Table of strains and plasmids

##### 2.1 Gene Sequences

**Table S2.** Genes used in this study. The B1PelB tag[2] is highlighted in blue, $His_6$ tag in pink and the thrombin cleavage site in grey. Start codons are highlighted in yellow and stop codons in red.

| Gene(s) | Nucleotide sequence (5' to 3') | Reference |
|---|---|---|
| **Curli: *csgB, csgA, csgG, csgE, csgF*** | ATGAAAAACAAATTGTTATTTATGATGTTAACAATACTGGGTGCGCCTGGGATTGCAGCCGCAGCAGG<br>TTATGATTTAGCTAATTCAGAATATAACTTCGCGGTAAATGAATTGAGTAAGTCTTCATTTAATCAGGC<br>AGCCATAATTGGTCAAGCTGGGACTAATAATAGTGCTCAGTTACGGCAGGGAGGCTCAAAACTTTTG<br>GCGGTTGTTGCGCAAGAAGGTAGTAGCAACCGGGCAAAGATTGACCAGACAGGAGATTATAACCTTG<br>CATATATTGATCAGGCGGGCAGTGCCAACGATGCCAGTATTTCGCAAGGTGCTTATGGTAATACTGCG | This study |

| | | |
|---|---|---|
| | ATGATTATCCAGAAAGGTTCTGGTAATAAAGCAAATATTACACAGTATGGTACTCAAAAAACGGCAAT<br>TGTAGTGCAGAGACAGTCGCAAATGGCTATTCGCGTGACACAACGTTAATTTCCATTCGACTTTTAAAT<br>CAATCCGATGGGGGTTTTACATGAAACTTTTAAAAGTAGCAGCAATTGCAGCAATCGTATTCTCCGGT<br>AGCGCTCTGGCAGGTGTTGTTCCTCAGTACGGCGGCGGCGGTAACCACGGTGGTGGCGGTAATAATA<br>GCGGCCCAAATTCTGAGCTGAACATTTACCAGTACGGTGGCGGTAACTCTGCACTTGCTCTGCAAACT<br>GATGCCCGTAACTCTGACTTGACTATTACCCAGCATGGCGGCGGTAATGGTGCAGATGTTGGTCAGG<br>GCTCAGATGACAGCTCAATCGATCTGACCCAACGTGGCTTCGGTAACAGCGCTACTCTTGATCAGTGG<br>AACGGCAAAAATTCTGAAATGACGGTTAAACAGTTCGGTGGTGGCAACGGTGCTGCAGTTGACCAGA<br>CTGCATCTAACTCCTCCGTCAACGTGACTCAGGTTGGCTTTGGTAACAACGCGACCGCTCATCAGTACT<br>AAtactagagccaggcataaagaggagaaatagtacaATGAAACGTTATTTACGCTGGATTGTGGCGGCAGAATT<br>TCTGTTCGCCGCAGGGAATCTTCACGCCGTTGAGGTAGAAGTCCCGGGATTGCTAACTGACCATACTG<br>TTTCATCTATTGGCCATGATTTTTACCGAGCCTTTAGTGATAAATGGGAAAGTGACTATACGGGTAACT<br>TAACGATTAATGAAAGGCCCAGTGCACGATGGGGAAGCTGGATCACTATAACGGTCAATCAGGACGT<br>TATTTTCCAGACTTTTTTATTTCCGTTGAAAAGAGACTTCGAGAAAACTGTCGTCTTTGCACTGATTCAA<br>ACTGAAGAAGCACTAAATCGTCGCCAGATAAATCAGGCGTTATTAAGTACGGGCGATTTGGCGCATG<br>ATGAATTCTAAATAAAAAATTGTTCGGAGGCTGCAATGCGTGTCAAACATGCAGTAGTTCTACTCATG<br>CTTATTTCGCCATTAAGTTGGGCTGGAACCATGACTTTCCAGTTCCGTAATCCAAACTTTGGTGGTAAC<br>CCAAATAATGGCGCTTTTTTATTAAATAGCGCTCAGGCCCAAAACTCTTATAAAGATCCGAGCTATAAC<br>GATGACTTTGGTATTGAAACACCCTCAGCGTTAGATAACTTTACTCAGGCCATCCAGTCACAAATTTTA<br>GGTGGGCTACTGTCGAATATTAATACCGGTAAACCGGGCCGCATGGTGACCAACGATTATATTGTCGA<br>TATTGCCAACCGCGATGGTCAATTGCAGTTGAACGTGACAGATCGTAAAACCGGACAAACCTCGACCA<br>TCCAGGTTTCGGGTTTACAAAATAACTCAACCGATTTTTAAGCCCCAGCTTCATAAGGAAAATAATCAT<br>GCAGCGCTTATTTCTTTTGGTTGCCGTCATGTTACTGAGCGGATGCTTAACCGCCCCGCCTAAAGAAGC<br>CGCCAGACCGACATTAATGCCTCGTGCTCAGAGCTACAAAGATTTGACCCATCTGCCAGCGCCGACGG<br>GTAAAATCTTTGTTTCGGTATACAACATTCAGGACGAAACCGGGCAATTTAAACCCTACCCGGCAAGT<br>AACTTCTCCACTGCTGTTCCGCAAAGCGCCACGGCAATGCTGGTCACGGCACTGAAAGATTCTCGCTG<br>GTTTATACCGCTGGAGCGCCAGGGCTTACAAAACCTGCTTAACGAGCGCAAGATTATTCGTGCGGCAC<br>AAGAAAACGGCACGGTTGCCATTAATAACCGAATCCCGCTGCAATCTTTAACGGCGGCAAATATCATG<br>GTTGAAGGTTCGATTATCGGTTATGAAAGCAACGTCAAATCTGGCGGGGTTGGGGCAAGATATTTTG<br>GCATCGGTGCCGACACGCAATACCAGCTCGATCAGATTGCCGTGAACCTGCGCGTCGTCAATGTGAGT<br>ACCGGCGAGATCCTTTCTTCGGTGAACACCAGTAAGACGATACTTTCCTATGAAGTTCAGGCCGGGGT<br>TTTCCGCTTTATTGACTACCAGCGCTTGCTTGAAGGGGAAGTGGGTTACACCTCGAACGAACCTGTTAT<br>GCTGTGCCTGATGTCGGCTATCGAAACAGGGGTCATTTTCCTGATTAATGATGGTATCGACCGTGGTC<br>TGTGGGATTTGCAAAATAAAGCAGAACGGCAGAATGACATTCTGGTGAAATACCGCCATATGTcggttc<br>caccggaatcctga | |
| ***Is*PETase$^{S238F}$<br>/W159H** | ATGGCCAATTTTCCGCGTGCAAGCCGTCTGATGCAGGCAGCAGTTTTAGGTGGTCTGATGGCAGTTAG<br>CGCAGCAGCAACCGCACAGACCAATCCGTATGCACGTGGTCCGAATCCGACCGCAGCAAGCCTGGAA<br>GCAAGCGCAGGTCCGTTTACCGTTCGTAGCTTTACCGTTAGCCGTCCGAGCGGTTATGGTGCAGGCAC<br>CGTTTATTATCCGACCAATGCCGGTGGCACCGTTGGTGCAATTGCCATTGTTCCGGGTTATACCGCACG<br>TCAGAGCAGCATTAAATGGTGGGGTCCGCGTCTGGCAAGTCATGGTTTTGTTGTTATTACCATTGATA<br>CCAACAGCACCCTGGATCAGCCGAGCAGCCGTAGCAGTCAGCAGATGGCAGCACTGCGTCAGGTTGC<br>CAGCCTGAATGGCACCAGCAGCAGCCCGATTTATGGTAAAGTTGATACAGCACGTATGGGTGTTATG<br>GGTCATAGCATGGGTGGTGGTGGTAGCCTGATTAGTGCAGCAAATAATCCGAGCCTGAAAGCAGCCG<br>CACCGCAGGCTCCGTGGGATAGCAGCACCAATTTTAGCAGCGTTACCGTTCCGACACTGATTTTTGCA<br>TGTGAAAATGATAGCATTGCACCGGTTAATAGCAGCGCACTGCCGATCTATGATAGTATGAGCCGTAA<br>TGCAAAACAGTTTCTGGAAATTAATGGTGGCAGCCATTTTTGTGCAAATAGCGGTAATAGCAATCAGG<br>CACTGATCGGTAAAAAGGGTGTTGCATGGATGAAACGCTTTATGGATAATGATACCCGCTATAGCACC<br>TTTGCCTGCGAAAATCCGAATAGCACCCGTGTTAGCGATTTTCGTACCGCAAATTGTAGCTAA | [3] |
| **PHL7** | ATGGCCAATCCGTATGAACGTGGTCCGGATCCGACCGAAAGCAGCATTGAAGCAGTTCGTGGTCCGT<br>TTGCAGTTGCACAGACCACCGTTAGCCGTCTGCAGGCAGATGGTTTTGGTGGTGGCACCATCTATTAT<br>CCGACAGATACCAGCCAGGGCACCTTTGGTGCAGTTGCCATTAGTCCGGGTTTTACCGCAGGTCAAGA<br>AAGCATTGCATGGCTGGGTCCGCGTATTGCAAGCCAGGGTTTTGTTGTTATTACCATTGATACCATTAC<br>GCGTCTGGATCAGCCGGATAGCCGTGGTCGTCAGCTGCAAGCAGCACTGGATCATCTGCGTACCAAT<br>AGCGTTGTTCGTAATCGTATTGATCCGAATCGTATGGCAGTTATGGGTCATAGCATGGGTGGTGGCG<br>GTGCACTGAGCGCAGCAGCAAATAATACCAGCCTGGAAGCAGCAATTCCGCTGCAAGGTTGGCATAC<br>CCGTAAAAATTGGAGCAGCGTTCGTACCCCGACACTGGTTGTTGGTGCACAGCTGGATACCATTGCAC<br>CGGTTAGCAGCCATAGCGAAGCCTTTTATAACAGCCTGCCGAGCGATCTGGATAAAGCATATATGGA<br>ACTGCGTGGTGCAAGCCATCTGGTTAGCAATACACCGGATACCACCACCGCAAAATATAGTATTGCCT<br>GGCTGAAACGTTTTGTGGATGATGATCTGCGTTATGAACAGTTTCTGTGTCCGGCACCGGATGATTTT<br>GCAATTAGCGAATATCGTAGCACGTGCCCGTTTTAA | [4] |
| ***Is*PETase$^{S238F}$<br>/W159H-His** | ATGGAACGTGCATGTGTTGCCGTTATGAAATATCTGCTGCCGACCGCAGCAGCGGGTCTGCTGCTGCT<br>GGCAGCACAACCTGCCATGGCCAATTTTCCGCGTGCAAGCCGTCTGATGCAGGCAGCAGTTTTAGGTG<br>GTCTGATGGCAGTTAGCGCAGCAGCAACCGCACAGACCAATCCGTATGCACGTGGTCCGAATCCGAC | This study |

| | | |
|---|---|---|
| | CGCAGCAAGCCTGGAAGCAAGCGCAGGTCCGTTTACCGTTCGTAGCTTTACCGTTAGCCGTCCGAGCG<br>GTTATGGTGCAGGCACCGTTTATTATCCGACCAATGCCGGTGGCACCGTTGGTGCAATTGCCATTGTT<br>CCGGGTTATACCGCACGTCAGAGCAGCATTAAATGGTGGGGTCCGCGTCTGGCAAGTCATGGTTTTGT<br>TGTTATTACCATTGATACCAACAGCACCCTGGATCAGCCGAGCAGCCGTAGCAGTCAGCAGATGGCA<br>GCACTGCGTCAGGTTGCCAGCCTGAATGGCACCAGCAGCAGCCCGATTTATGGTAAAGTTGATACAG<br>CACGTATGGGTGTTATGGGTCATAGCATGGGTGGTGGTGGTAGCCTGATTAGTGCAGCAAATAATCC<br>GAGCCTGAAAGCAGCCGCACCGCAGGCTCCGTGGGATAGCAGCACCAATTTTAGCAGCGTTACCGTT<br>CCGACACTGATTTTTGCATGTGAAAATGATAGCATTGCACCGGTTAATAGCAGCGCACTGCCGATCTA<br>TGATAGTATGAGCCGTAATGCAAAACAGTTTCTGGAAATTAATGGTGGCAGCCATTTTTGTGCAAATA<br>GCGGTAATAGCAATCAGGCACTGATCGGTAAAAAGGGTGTTGCATGGATGAAACGCTTTATGGATAA<br>TGATACCCGCTATAGCACCTTTGCCTGCGAAAATCCGAATAGCACCCGTGTTAGCGATTTTCGTACCGC<br>AAATTGTAGCCTGGTTCCGCGTGGTAGTcatcatcatcatcatcacTAA | |
| **PHL7-His** | ATGGAACGTGCATGTGTTGCCGTTATGAAATATCTGCTGCCGACCGCAGCAGCGGGTCTGCTGCTGCT<br>GGCAGCACAACCTGCCATGGCCAATCCGTATGAACGTGGTCCGGATCCGACCGAAAGCAGCATTGAA<br>GCAGTTCGTGGTCCGTTTGCAGTTGCACAGACCACCGTTAGCCGTCTGCAGGCAGATGGTTTTGGTGG<br>TGGCACCATCTATTATCCGACAGATACCAGCCAGGGCACCTTTGGTGCAGTTGCCATTAGTCCGGGTT<br>TTACCGCAGGTCAAGAAAGCATTGCATGGCTGGGTCCGCGTATTGCAAGCCAGGGTTTTGTTGTTATT<br>ACCATTGATACCATTACGCGTCTGGATCAGCCGGATAGCCGTGGTCGTCAGCTGCAAGCAGCACTGG<br>ATCATCTGCGTACCAATAGCGTTGTTCGTAATCGTATTGATCCGAATCGTATGGCAGTTATGGGTCATA<br>GCATGGGTGGTGGCGGTGCACTGAGCGCAGCAGCAAATAATACCAGCCTGGAAGCAGCAATTCCGCT<br>GCAAGGTTGGCATACCCGTAAAAATTGGAGCAGCGTTCGTACCCCGACACTGGTTGTTGGTGCACAG<br>CTGGATACCATTGCACCGGTTAGCAGCCATAGCGAAGCCTTTTATAACAGCCTGCCGAGCGATCTGGA<br>TAAAGCATATATGGAACTGCGTGGTGCAAGCCATCTGGTTAGCAATACACCGGATACCACCACCGCAA<br>AATATAGTATTGCCTGGCTGAAACGTTTTGTGGATGATGATCTGCGTTATGAACAGTTTCTGTGTCCG<br>GCACCGGATGATTTTGCAATTAGCGAATATCGTAGCACGTGCCCGTTTCTGGTTCCGCGTGGTAGTcat<br>catcatcatcatcacTAA | This study |

#### 2.2 Plasmid assembly

pAg43 (pBAD-Ag43) was purchased from Addgene (Addgene plasmid #107743; http://n2t.net/addgene:107743; RRID: Addgene 107743). pJUMP29-1D(sfGFP) was a gift from Prof. Chris French (University of Edinburgh) and is also available through Addgene (Addgene plasmid # 126981; http://n2t.net/addgene:126981; RRID: Addgene 126981).

PCR amplified and purified genes PHL7 and *Is*PETase$^{S238F/W159H}$ were cloned into a pET22b-PelB1 backbone using NdeI and XhoI restriction sites and ligated with T4 ligation using the protocol previously mentioned. The genes for B1PelB-*Is*PETase$^{S238F/W159H}$ and B1PelB-PHL7 were amplified out of the pET22b backbone using the primers shown in Table S3. The B1PelB-*Is*PETase$^{S238F/W159H}$ and B1PelB-PHL7 were assembled using BsaI-HF®v2 (NEB) mediated cleavage and T4 ligase for the ligation into a pET28a, resulting in C-terminal thrombin cleavage site and a $His_6$-tag being added to B1PelB-*Is*PETase$^{S238F/W159H}$ and B1PelB-PHL7. A plasmid was assembled using JUMP modular cloning[5] into a pJUMP29-1D plasmid. The gene was assembled under the control of a Ptac promotor with RBS pET. The pJUMP backbone, B1PelB-*Is*PETase$^{S238F/W159H}$-$His_6$ and B1PelB-PHL7-$His_6$ were amplified using the primers in Table S3 and assembled using BsaI-HF®v2 and T4 ligase to generate the plasmids _pPHL7 and _p*Is*PETase (Table S4) used in this study.

pCurli was assembled by amplification of *csgBA* and *csgGEF* from the genome of *E. coli* MG1655 using primer pairs csgG fwd/csgA rev and csgG fwd/csgF rev (Table S3). The assembled operon and pAg43 backbone were amplified using the primer pairs (Table S3) and assembled using BsmBI-v2 mediated cleavage and T4 ligase. To produce pBAD (no adhesion control), primers were used to amplify pAg43 backbone and then subsequently digested with BsmBI-v2 and re-ligated with T4 ligase.

**Table S3.** Oligos used in this study to assemble plasmids. Enzyme recognition sites are underlined, while overhangs are labelled in blue if relevant.

| Oligo name | Primer (5'…3') | Purpose |
|---|---|---|
| **BsaI_PelB_fwd** | ggctacggtctcgtggaacgtgcatgtgttgCCGTTATG | Amplification of PET hydrolase gene for insertion into pET28a |
| **BsaI_PHL7_rev** | ggctacggtctcggatgACTACCACGCGGAACCAGAAACGGGCACGTGCTACGATATTC | |
| **BsaI_PETase_rev** | ggctacggtctcggatgACTACCACGCGGAACCAGGCTACAATTTGCGGTACGAAAATCGC | |
| **BsaI_pET28_fwd** | ggctacggtctcgcatcatcatcatcatcacTAACATATGGCTAGCATGACTGGTGGAC | Amplification of pET28a backbone |
| **BsaI_pET28_rev** | ggctacggtctcgtccatggtatatctccttcttaaagttaaacaaaattatttc | |
| **BsaI_pET28a-PelB1_fwd** | ggctacggtctcgtactttgtttaactttaagaaggagatataccatggaacgtgcatgtgttgCC | Amplification of PET hydrolase gene for insertion into pJUMP29-1D |
| **BsaI_pET28a-thrombin_rev** | ggctacggtctcgAAGCTTAgtgatgatgatgatgatgACTACCACGCGGAACCAG | |
| **BsaI_JUMP29_fwd** | ggctacggtctcgGCTTtcgacccgcatgttcgcatg | Amplification of pJUMP backbone |
| **BsaI_JUMP29_rev** | ggctacggtctcgagtaaattgtgagcgctcacaattccacacattatacg | |
| **csgB fwd** | ctagagaaagaggagaaatactagaATGAAAAACAAATTGTTATTTATGATGTTAAC | Amplification of curli genes out of the *E. coli* genome |
| **csgA rev** | ctcctctttatgcctggctctagtaTTAGTACTGATGAGCGGTC | |
| **csgG fwd** | tactagagccaggcataaagaggagaaatagtacaATGAAACGTTATTTACGCTGG | |
| **csgF rev** | cgactgagcctttcgttttatttgTCAGGATTCCGGTGGAAC | |
| **BsmBI curli fwd** | ggctaccgtctcgactagaATGAAAAACAAATTGTTATTTATGATGTTAACAATACTGG | Amplification of csg operon for insertion into pBAD backbone |
| **BsmBI curli rev** | ggctaccgtctcgctctagtatcaggattccggtggaaccg | |
| **BsmBI pBAD fwd** | ggctaccgtctcgagagccaggcatcaaataaaacgaaaggc | Amplification of pBAD backbone for insertion of csg operon |
| **pBAD EV rev** | ggctaccgtctcatagtatttctcctctttctctagtagctagcccaaaaaaacg | |
| | ggctaccgtctcatagtatttctcctctttctctagtagctagcccaaaaaaacg | Amplification of pBAD backbone for empty vector |
| **pBAD EV fwd** | ggctaccgtctcaactagacaaataaaacgaaaggctcagtcgaaagactgg | |

**Table S4**. Plasmids used in this study.

| Plasmid name | Proteins encoded | Backbone | Promoter | Reference |
|---|---|---|---|---|
| **pBAD** | N/A | pBAD | araBAD | This study |
| **pCurli** | CsgB, CsgA, CsgG, CsgE, CsgF | pBAD | | |
| **pAg43** | Ag43 | | | [6] |
| **p*Is*PETase** | B1PelB- IsPETase$^{S238F/W159H}$-His$_6$ | pJUMP29-1D | Ptac | This study |
| **pPHL7** | B1PelB-PHL7-His$_6$ | | | |
| **pJUMP29-1D** | sfGFP | | BBa_J23100 | [5] |

#### 2.3 Strains

##### 2.3.1 Knockout strain generation

*E. coli* MG1655Δ*csgA*, MG1655Δ*csgA* and MG1655Δ*csgA*Δ*flu* strains were generated following the pCas/pTarget F protocol[7] with minor modifications[8], as follows. All N20s were identified using CHOPCHOP[9] screening for targets of *csgA* and *flu* in the *E. coli* MG1655. Two different N20s were chosen to target either the 5' or 3' end of either gene (Table S5). The N20s were screened against the *E. coli* str. K-12 substr. MG1655 (U000096.3) genome using BLAST[10] to determine there were no hits of >15 base-pair homology and the IDT OligoAnalyser tool[11] was used to screen for secondary structures. For the annealing of the N20s oligos, the reaction mix ratio was adjusted (43 µL water, 5 µL T4 ligase buffer, 1 µL each of 100 µM forward and reverse N20 oligo) and the PCR was cooled at 5 °C per minute to 25 °C. BsaI-HF®v2 (NEB) was used to digest pEcgRNA and the ~2 kb desired band was gel extracted. The gel-extracted pEcgRNA backbone and N20s were ligated using a T4 ligase.

For the synthesis of the donor DNA that will be used to replace the genes, the donor DNA for the knockout of *flu* comprised the flanking genomic region. Due to the highly repetitive region around the *csgA* gene, primers generated formed secondary structures and to overcome this the homology arms were shifted into *csgA* and a stop codon (TAG) was introduced so that only a tripeptide would be generated. Donor DNA was assembled by amplifying homology arm 1 and 2 separately using 1 µL boiled *E. coli* MG1655 as a genomic template (5 µL 10 mM dNTPs, 2.5 µL each forward and reverse oligo, 10 µL Q5 reaction buffer, 0.5 µL Q5 DNA polymerase, 1.5 µL DMSO, 27 µL MilliQ-water). Overlap extension PCR[12] was used to generate the donor DNA and then gel-purified. The gel-purified donor DNA was then reamplified using the H1 fwd/H2 rev primers (Table S5) and PCR purified to generate the final product.

Electro-competent *E. coli* MG1655 cells containing pEcCas plasmid (90 µL) were electroporated with 100 ng of pEcgRNA plasmid containing the N20s and 350-400 ng donor DNA in Gene Pulser/MicroPulser Electroporation Cuvettes (Bio-Rad). The plasmids were cured from successful knockouts as specified previously[7]. MG1655Δ*csgA* was used as the parent strain to generate the double knockout (Table S6).

**Table S5.** Oligos used to generate knockouts. The overhangs (blue) compatible with the BsaI-generated overhangs in pEcgRNA resulting of the insertion of the N20s into pEcgRNA.

| Oligo name | Primer (5'…3') | Purpose in generation of knockouts |
|---|---|---|
| **f_flu_5_N20** | TAGTGGTCTTCAGAGAGTGAACCCCGG | N20 for targeting CRISPR-Cas9 |
| **r_flu_5_N20** | AAACCCGGGGTTCACTCTCTGAAGACC | |
| **f_flu_3_N20** | TAGTACGTTCTCTCCCTCACAGAATGG | |
| **r_flu_3_N20** | AAACCCATTCTGTGAGGGAGAGAACGT | |
| **f_csgA_5_N20** | TAGTGGTGTTGTTCCTCAGTACGGCGG | |
| **r_csgA_5_N20** | AAACCCGCCGTACTGAGGAACAACACC | |
| **f_csgA_3_N20** | TAGTCCGCTCATCAGTACTAATACATC | |
| **r_csgA_3_N20** | AAACGATGTATTAGTACTGATGAGCGG | |
| **H1_flu_fwd** | CTGAGTCTGCTCACAAAAGCACTGTTTTCGTTA | Construction of donor DNA/homology arms |
| **H1_flu_rev** | GGCGATGGTTCTGTCACAGCTTTTCCTTAGATTGAGGG TGAATAAAAAAGCCGGTACCCACAAAC | |
| **H2_flu_fwd** | ATCTAAGGAAAAGCTGTGACAGAACCATCGCCTCTCTG TGGTCCCGG | |
| **H2_flu_rev** | TCCAGCGCGATATGTTTACCATCAAGTCCC | |
| **H1_csgA_fwd** | CAACATGAAAAACAAATTGTTATTTATG | |

| | |
|---|---|
| **H1_csgA_rev** | ACAAATGATGTACTAAAGTTTCATGTAAAACCCCCATCG GATTGATTT |
| **H2_csgA_fwd** | TACATGAAACTTTAGTACATCATTTGTATTACAGAAACA GGGCGC |
| **H2_csgA_rev** | TTCAACAAATTAAGACTTTTCTGAAGA |

**Table S6.** Bacterial strains used in this study.

| Strain name | Genotype | Antibiotic resistance |
|---|---|---|
| **MG1655Δ*flu*_pAg43** | *E. coli* MG1655Δ*flu* _pAg43 | Ampicillin |
| **MG1655Δ*flu*_pCurli** | *E. coli* MG1655Δ*flu* _pCurli | |
| **MG1655Δ*csgA*_pAg43** | *E. coli* MG1655Δ*csgA* _pAg43 | |
| **MG1655Δ*csgA*_pCurli** | *E. coli* MG1655Δ*csgA* _pCurli | |
| **MG1655Δ*csgA*Δ*flu*_pAg43** | *E. coli* MG1655Δ*csgA*Δ*flu* _pAg43 | |
| **MG1655Δ*csgA*Δ*flu*_pCurli** | *E. coli* MG1655Δ*csgA*Δ*flu* _pCurli | |
| **MG1655Δ*csgA*Δ*flu*_pBAD** | *E. coli* MG1655Δ*csgA*Δ*flu* Empty pBAD | |
| **MG1655Δ*csgA*Δ*flu*_pPHL7** | *E. coli* MG1655Δ*csgA*Δ*flu* pPHL7 | Kanamycin |
| **MG1655Δ*csgA*Δ*flu*_pIsPETase** | *E. coli* MG1655Δ*csgA*Δ*flu* pIsPETase$^{S238F/W159H}$ | |
| **MG1655Δ*csgA*Δ*flu*_pJUMP29-1D** | *E. coli* MG1655Δ*csgA*Δ*flu* pJUMP29-1D | |
| **MG1655Δ*csgA*Δ*flu*-_pAg43_pPHL7** | *E. coli* MG1655Δ*csgA*Δ*flu* pAg43, pPHL7 | Ampicillin, kanamycin |
| **MG1655Δ*csgA*Δ*flu*-_pCurli_pPHL7** | *E. coli* MG1655Δ*csgA*Δ*flu* pCurli, pPHL7 | |
| **MG1655Δ*csgA*Δ*flu*-_pBAD_pPHL7** | *E. coli* MG1655Δ*csgA*Δ*flu* pBAD, pPHL7 | |

### 3 Plastic substrates

**Table S7**. Plastics used throughout the study and their specifications. N/A: non-applicable.

| Plastic | Origin | Type | Code | Specifications |
|---|---|---|---|---|
| **Poly(ethylene terephthalate)** | DuPont Teijin films | Commercial | N/A | Melinex OD<br>Thickness: 125µm<br>Surface non-treated |
| | Goodfellow | Commercial | GF1545975 6-1EA | Thickness: 0.125mm<br>Structure: Biaxially Oriented |
| | A. Pickford Lab (University of Portsmouth) | Commercial | N/A | Cryo-milled (2 mM particle size) |
| **Poly(propylene)** | Goodfellow | Commercial | 185-275-23 | Thickness: 0.05mm<br>Structure: Biaxially Oriented |

| | | | | |
|---|---|---|---|---|
| **Poly(lactic acid)** | Goodfellow | Commercial | 700-387-99 | Thickness: 0.05mm<br>Structure: Amorphous |
| **Low-density poly(ethylene)** | Goodfellow | Commercial | 9002884 | Thickness: 0.05mm<br>Structure: Biaxially Oriented |
| **Poly(tetrafluoro ethyelene)** | Camthorne Ind Supplied Ltd | Commercial | G400 | Thickness: 1.5mm |
| **Poly(ethylene terephthalate)** | Unknown | Post-consumer | N/A | Bottle |
| **Poly(propylene)** | Haribo | Post-consumer | N/A | Bag |
| **Poly(lactic acid)** | Vegware | Post-consumer | N/A | Cup |
| **Low-density poly(ethylene)** | Unknown | Post-consumer | N/A | Packaging |

### 4 Additional experimental data

##### 4.1.1 Media recipe

**Table S8**. Recipe for M63 media[13] containing glucose and casamino acids[6].

| Component | | Final concentration |
|---|---|---|
| **5x M63 media** | Potassium dihydrogen phosphate | 100 mM |
| | Ammonium sulphate | 15 M |
| **Additives** | Magnesium sulphate | 1 mM |
| | Iron (III) hexahydrate | 0.0018 mM |
| | Succinic acid disodium hexahydrate | 17 mM |
| | Casamino acids | 0.1% |
| | Glucose | 11.1 mM (0.2%) |

##### 4.1.2 Crystal violet staining and ImageJ analysis of adhesion to PS plates

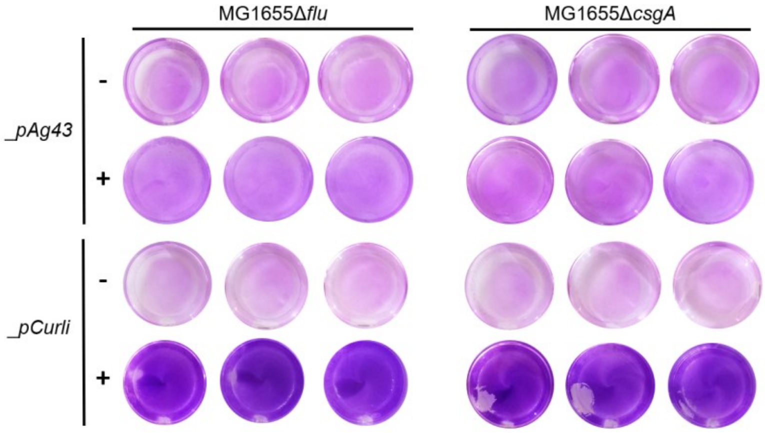

**Figure S1**. Inducible adhesion of *E. coli* single knockouts to PS surfaces, images of stained PS-plate adhered cells showing biomass distribution. Uninduced: - ; induced: + . Conditions: 0.1% arabinose, 30 °C, 230 rpm, 20 h.

**Table S9.** Statistical analysis (max, min, intensity of mode, count) of grayscale histograms generated by ImageJ of cell-adhered PS wells (Figure 2a, Figure S1).

| Strain | Plasmid | Induced | Replicate number | Count | Intensity of mode | Max | Min | StdDev |
|---|---|---|---|---|---|---|---|---|
| **MG1655** | pAg43 | - | 1 | 22711 | 1172 | 255 | 136 | 14.168 |
| | | | 2 | 22703 | 1056 | 248 | 177 | 10.503 |
| | | | 3 | 23243 | 1028 | 240 | 145 | 10.231 |
| | | + | 1 | 21396 | 605 | 243 | 149 | 16.486 |
| | | | 2 | 20224 | 847 | 255 | 149 | 13.062 |
| | | | 3 | 20224 | 1135 | 229 | 133 | 11.481 |
| | pCurli | - | 1 | 24194 | 740 | 254 | 151 | 13.837 |
| | | | 2 | 24194 | 1069 | 255 | 158 | 10.663 |
| | | | 3 | 24742 | 1203* | 253 | 161 | 9.835 |
| | | + | 1 | 21002 | 350 | 242 | 88 | 32.893 |
| | | | 2 | 19480 | 272 | 252 | 85 | 36.602 |
| | | | 3 | 21111 | 382 | 238 | 91 | 28.964 |
| **MG1655Δ*flu*** | pAg43 | - | 1 | 24208 | 791 | 246 | 148 | 16.311 |
| | | | 2 | 25876 | 923 | 255 | 150 | 12.346 |
| | | | 3 | 26876 | 1424 | 254 | 143 | 9.614 |
| | | + | 1 | 26596 | 1667 | 250 | 153 | 9.162 |
| | | | 2 | 27032 | 1697 | 249 | 153 | 8.526 |
| | | | 3 | 25308 | 1228 | 253 | 142 | 10.504 |
| | pCurli | - | 1 | 23658 | 843 | 248 | 166 | 14.121 |
| | | | 2 | 24601 | 1014 | 254 | 165 | 12.938 |
| | | | 3 | 23382 | 999 | 254 | 162 | 11.020 |
| | | + | 1 | 23784 | 640 | 247 | 79 | 25.623 |
| | | | 2 | 23644 | 636 | 225 | 76 | 20.006 |
| | | | 3 | 23644 | 761 | 235 | 91 | 18.638 |
| **MG1655Δ*csgA*** | pAg43 | - | 1 | 27900 | 1144 | 230 | 143 | 15.169 |
| | | | 2 | 26724 | 1257 | 253 | 151 | 13.266 |
| | | | 3 | 26724 | 1489 | 255 | 163 | 11.881 |
| | | + | 1 | 26024 | 931 | 255 | 139 | 14.832 |
| | | | 2 | 25168 | 1096 | 255 | 142 | 12.608 |
| | | | 3 | 24053 | 1082 | 254 | 155 | 11.428 |
| | pCurli | - | 1 | 28796 | 1309 | 255 | 150 | 12.600 |

| | | | | | | | | |
|---|---|---|---|---|---|---|---|---|
| | | | 2 | 28360 | 1182 | 253 | 169 | 12.571 |
| | | | 3 | 24736 | 1523 | 253 | 151 | 10.691 |
| | | + | 1 | 24194 | 603 | 249 | 59 | 27.053 |
| | | | 2 | 24328 | 477 | 204 | 83 | 24.010 |
| | | | 3 | 22445 | 596 | 219 | 88 | 19.818 |
| **MG1655Δ*csgA*Δ*flu*** | pAg43 | - | 1 | 22960 | 700 | 253 | 137 | 16.996 |
| | | | 2 | 22576 | 1002 | 249 | 167 | 10.535 |
| | | | 3 | 23240 | 1121 | 251 | 161 | 12.353 |
| | | + | 1 | 22574 | 715 | 246 | 130 | 15.563 |
| | | | 2 | 24052 | 1248 | 254 | 156 | 10.190 |
| | | | 3 | 23780 | 1244 | 255 | 142 | 13.240 |
| | pCurli | - | 1 | 23659 | 1344 | 248 | 176 | 9.859 |
| | | | 2 | 23382 | 1447 | 240 | 169 | 8.536 |
| | | | 3 | 22286 | 1205 | 248 | 177 | 9.042 |
| | | + | 1 | 23092 | 619 | 255 | 78 | 61.345 |
| | | | 2 | 23520 | 557 | 252 | 73 | 65.157 |
| | | | 3 | 24752 | 668 | 252 | 69 | 56.849 |

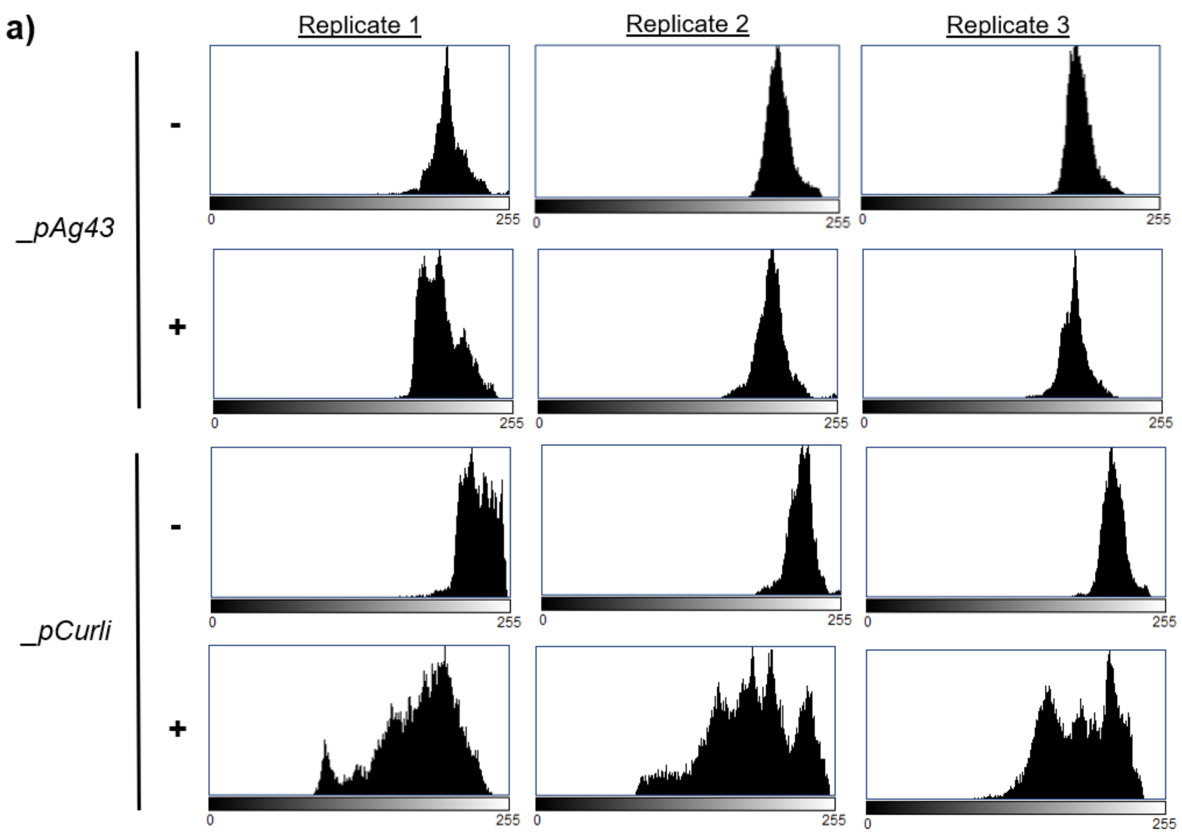

b)

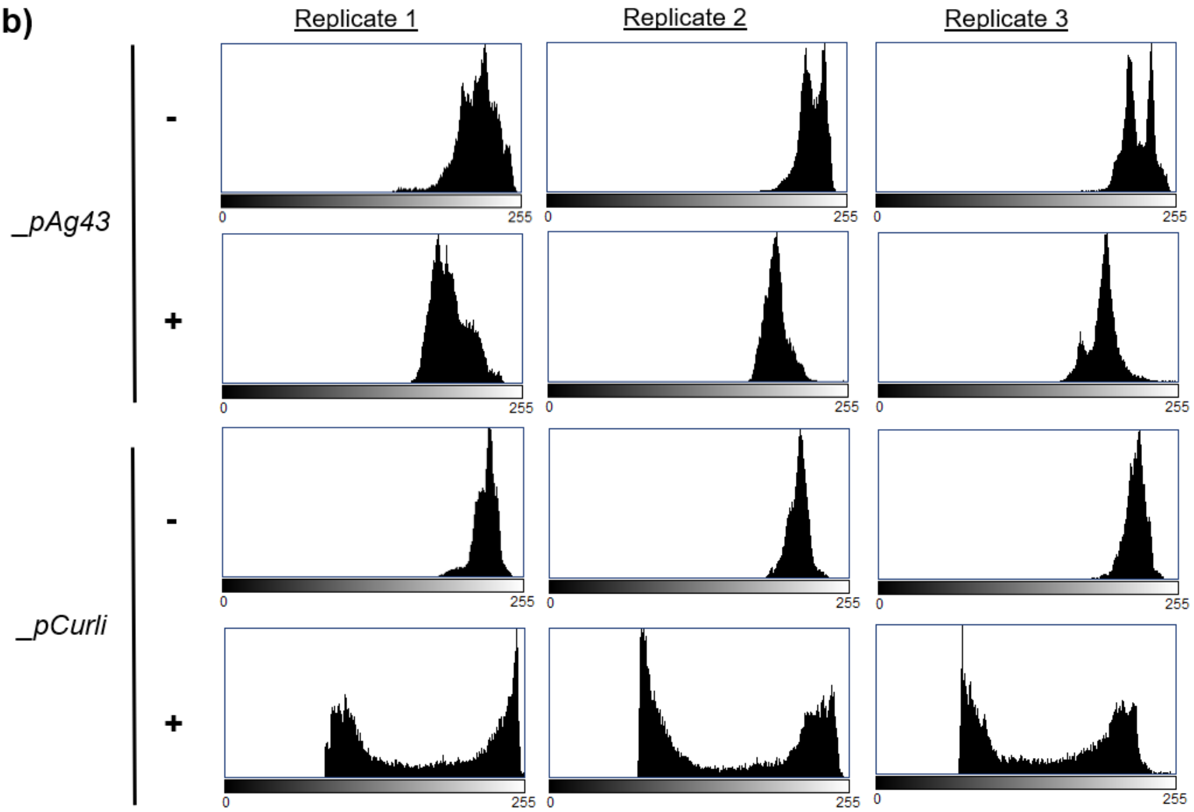
Replicate 1
Replicate 2
Replicate 3
_pAg43
_pCurli
-
+
0
255

c)

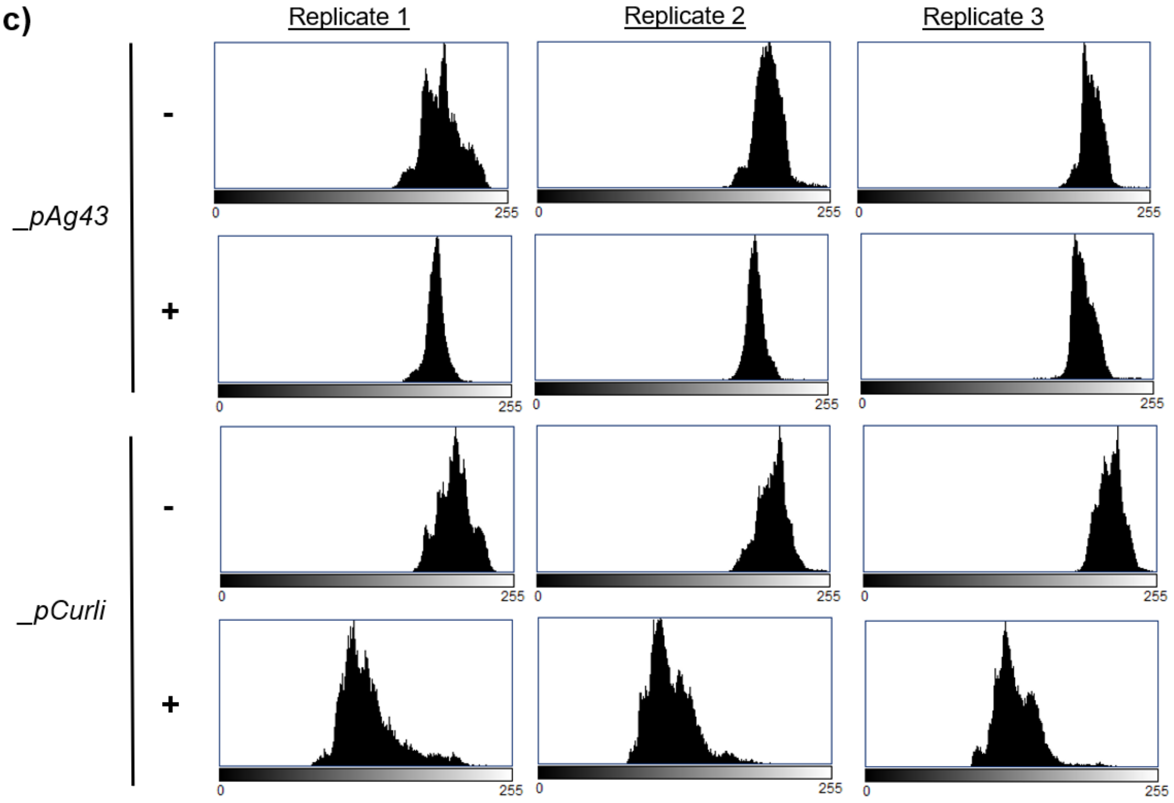
Replicate 1
Replicate 2
Replicate 3
_pAg43
_pCurli
-
+
0
255

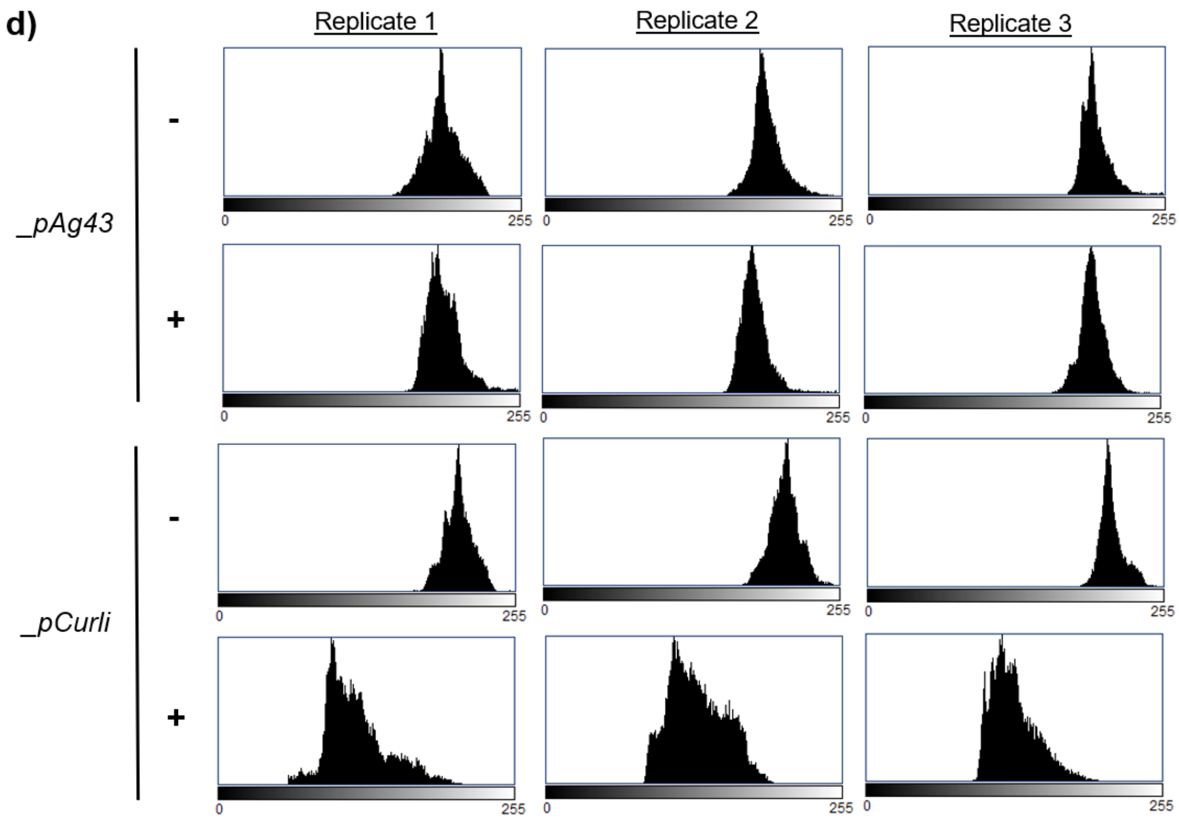

**Figure S2.** Grayscale histograms generated by ImageJ of cell-adhered PS wells (Figure 2b) for **(a)** MG1655, **(b)** MG1655Δ*csgA*Δ*flu,* **(c)** MG1655Δ*flu* and **(d)** MG1655Δ*csgA* showing distribution and intensity of coverage of PS wells.

##### 4.1.3 Engineered adhesion to post-consumer plastics

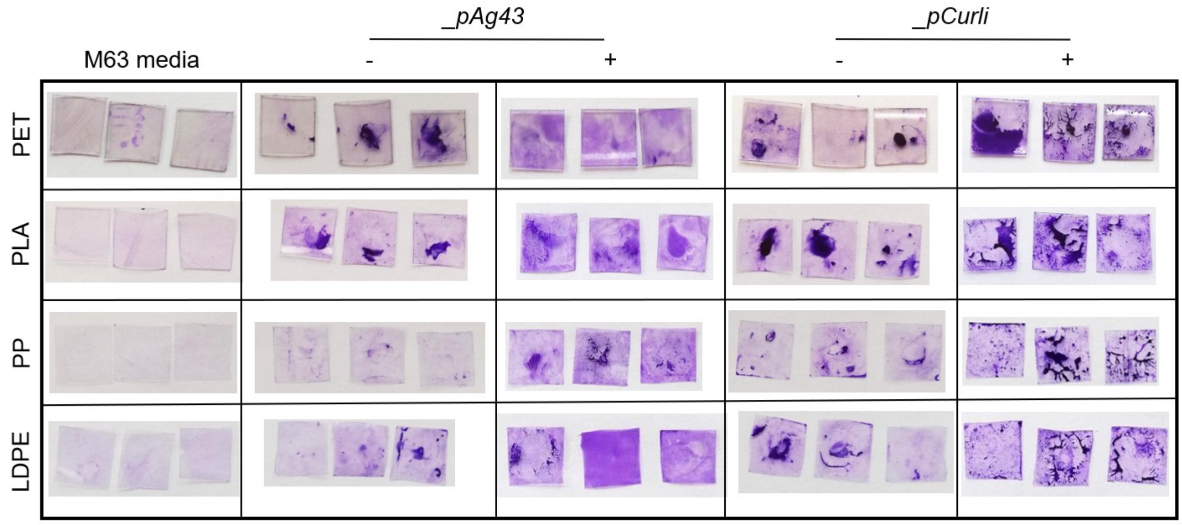

**Figure S3**. Adhesion of *E. coli* MG1655Δ*csgA*Δ*flu* expressing _pAg43 or _pCurli to different commercial plastics. Crystal violet imaging showing biomass distribution across the post-consumer plastic surface. Uninduced: - ; induced: + . Conditions: 0.1% arabinose, 30 °C, 230 rpm, 20 h.

##### 4.1.4 Water contact angle measurements

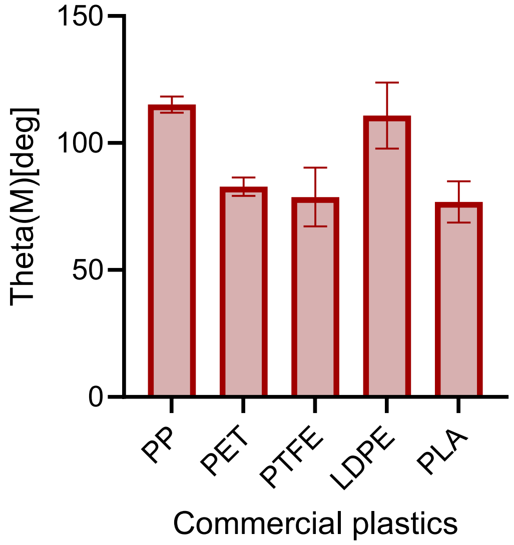

**Figure S4**. Water contact angle measurement using a 500 µL Hamilton syringe and a Drop Shape Analyser DSA25.

##### 4.1.5 Expression and activity of PET hydrolases

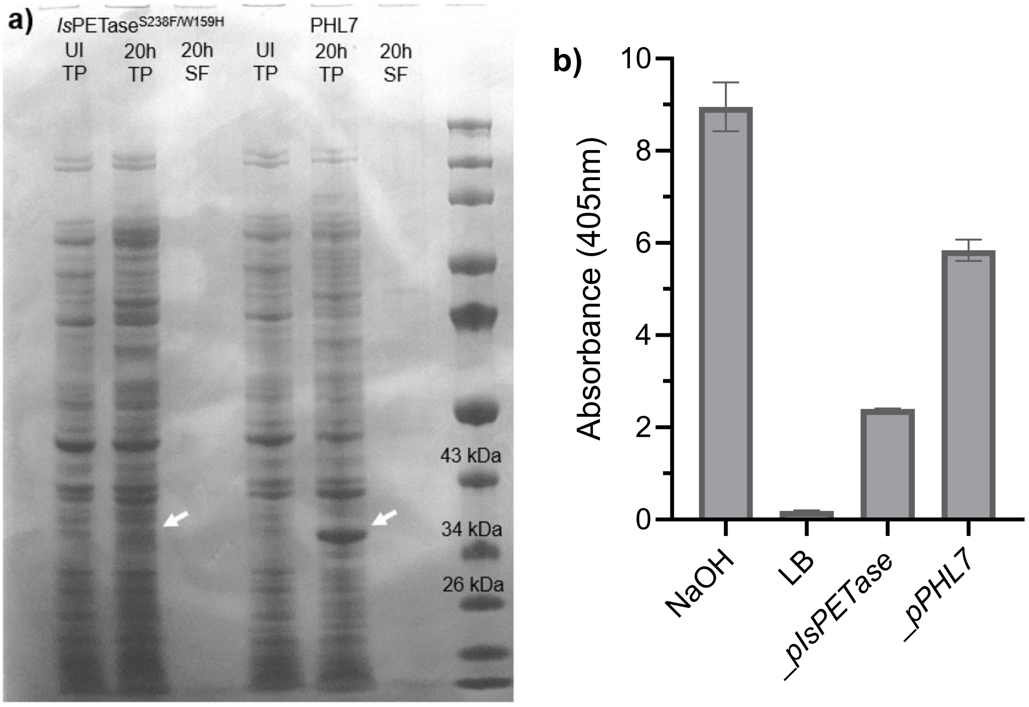

**Figure S5**. SDS-PAGE for uninduced (UI) and end-point (20h) expression of *Is*PETase$^{S238F/W159H}$ and PHL7 **(a)** pJUMP29 in MG1655Δ*csgA*Δ*flu.* Conditions: LB media, final 0.45 mM IPTG, 0.1% arabinose, 20 °C. **(b)** *p*-nitrophenyl butyrate assay on secreted fraction of *Is*PETase$^{S238F/W159H}$ and PHL7 in MG1655Δ*csga*Δ*flu*. Conditions: LB media, final 0.45 mM IPTG, 20 °C. 0.25 M NaOH positive control, final concentration 12.5 mM. Colour Protein Ladder (NEB). TP: total protein; SF: secreted fraction.

##### 4.1.6 Optimisation of pairing adhesion and secretion

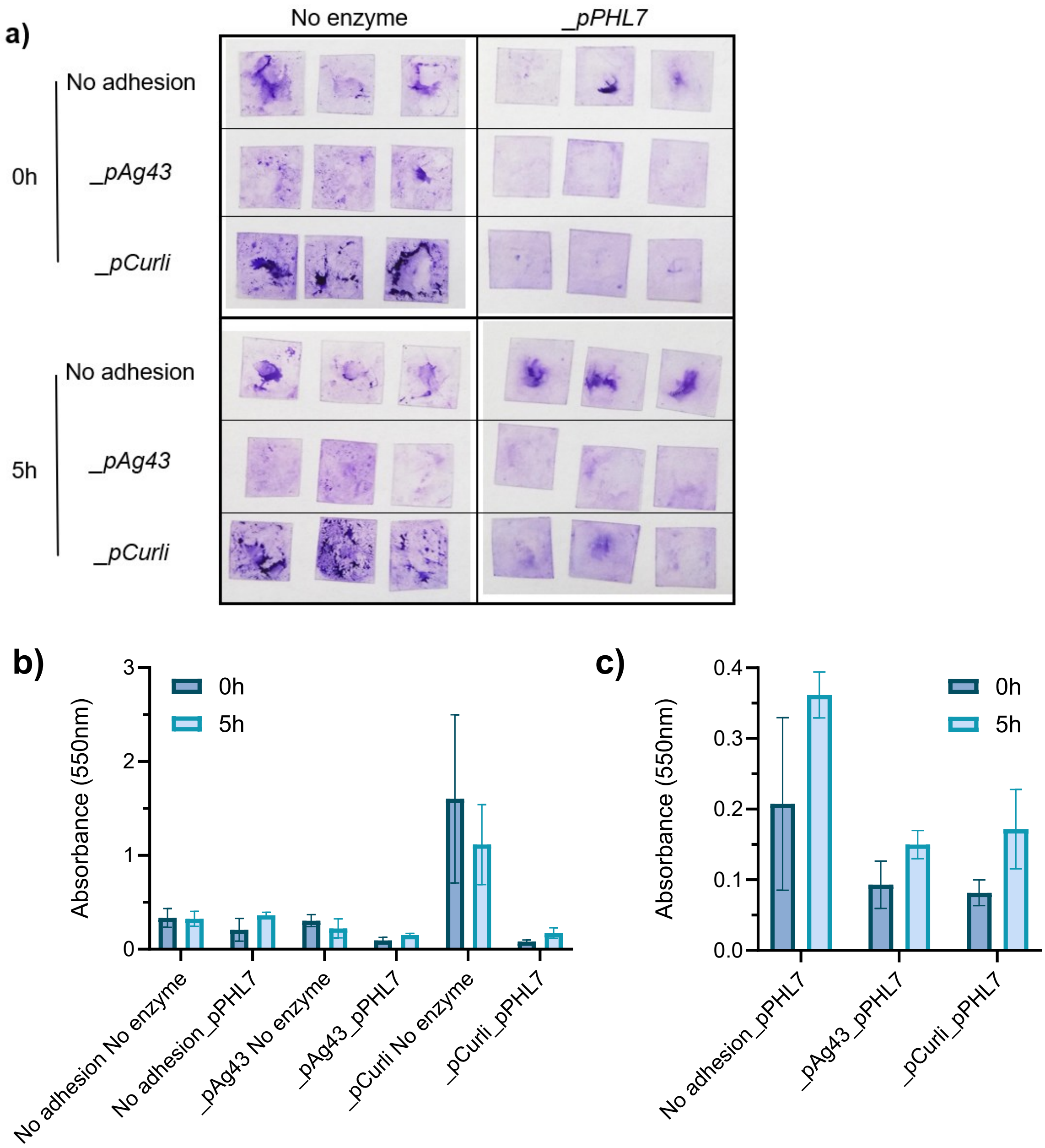

**Figure S6.** Optimisation of pairing of adhesion and secretion in MG1655Δ*csgA*Δ*flu* investigating a 0h and 5h time gap between the induction of adhesion (arabinose) and secretion (final 0.45 mM IPTG). Crystal violet **(a)** imaging to determine biofilm distribution across PET plastic surface and **(b)** quantification of adhesion as well as a **(c)** focus on the strains with _pPHL7 (same data set). Conditions: 0.1% arabinose, M63 media, 30 °C, 230 rpm, 22 h.

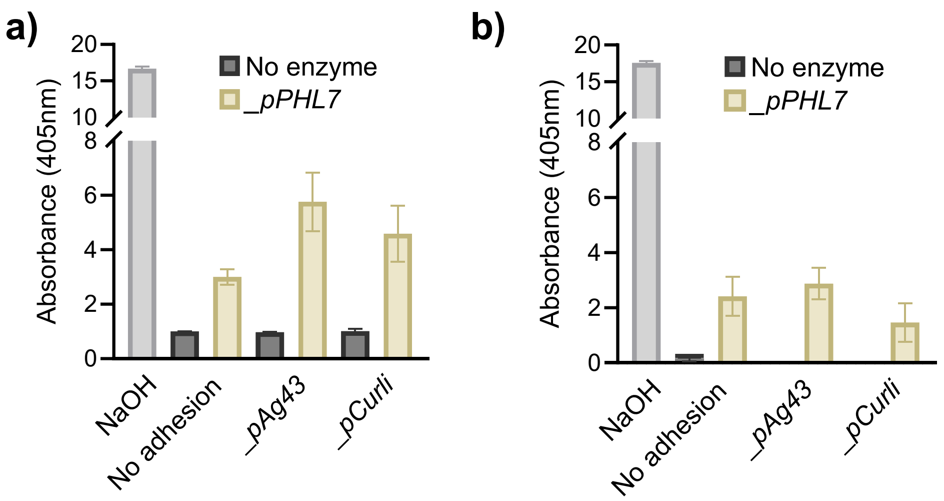

**Figure S7.** Optimisation of pairing adhesion with secretion MG1655Δ*csgA*Δ*flu* investigating activity of PHL7 using a pNPB assay with either final **(a)** 0.1 mM or **(b)** 0.45 mM IPTG for induction. 0.25 mM NaOH positive control, final concentration 12.5 mM. Conditions: 0.1% arabinose, M63 media, 5 h gap between inductions.
